# Chloroplast redox state changes indicate cell-to-cell signalling during the hypersensitive response

**DOI:** 10.1101/2021.02.08.430316

**Authors:** Tjaša Lukan, Anže Županič, Tjaša Mahkovec Povalej, Jacob O. Brunkard, Mojca Juteršek, Špela Baebler, Kristina Gruden

**Affiliations:** National Institute of Biology, Večna pot 111, 1000 Ljubljana, Slovenia; Laboratory of Genetics, University of Wisconsin – Madison, Madison, WI 53706, USA

**Keywords:** *Solanum tuberosum* (potato), virus resistance, *Potyvirus*, hypersensitive response (HR)-conferred resistance, salicylic acid, programmed cell death, chloroplastic reactive oxygen species, chloroplast redox state, spatiotemporal analysis, stromules, immune signalling

## Abstract

Hypersensitive response (HR)-conferred resistance is associated with an induction of programmed cell death (PCD) and pathogen spread restriction in its proximity. While the pivotal role of salicylic acid (SA) in the restriction of pathogen spread during HR has been confirmed, the exact role of chloroplastic reactive oxygen species and the link between SA signalling and chloroplast redox state during HR remain unexplained. To unravel these relationships, we performed detailed spatiotemporal analysis of chloroplast redox response to potato virus Y (PVY) infection in resistant *Ny-1*-gene-bearing potato and its transgenic counterpart with impaired SA accumulation and compromised resistance. We found that the chloroplasts are highly oxidized in the cells adjacent to the cell death zone at different stages after virus inoculation in both genotypes. Moreover, we detected individual cells with moderately oxidized chloroplasts, which we call “signalling cells”, in close proximity as well as farther from the cell death zone. These are relatively rare in SA-deficient plants, suggesting their role in signalling for HR-conferred resistance. This hypothesis is further supported by highly induced formation of stroma filled tubules that extend from chloroplasts (stromules) in the cells adjacent to signalling cells. Unexpectedly, the cells with the highest occurrence of stromules have chloroplasts in more reduced state than the adjacent ones. In addition, we show that stromules are induced also at the edge of PVY multiplication zone. We conclude that chloroplast redox state and stromule formation are tightly spatiotemporally regulated by SA-signalling which leads to effective HR response.

## Introduction

Plants have evolved a range of constitutive and inducible resistance mechanisms to respond to pathogen attack. The more specific layers of immunity are mediated by intracellular resistance (R) proteins (Jones & Dangl, 2006), which confer the recognition of pathogen-derived effectors and initiate effector-triggered immunity (ETI). Successful ETI often results in hypersensitive response (HR)-conferred resistance, where restriction of pathogens to the infection site is associated with localized programmed cell death (PCD) (Künstler *et al.*, 2016). HR is preceded by a series of biochemical and cellular signals, including salicylic acid (SA) biosynthesis in chloroplasts and production of reactive oxygen species (ROS) in chloroplasts and apoplast (Lu & Yao, 2018; Balint-Kurti, 2019). Besides hosting the biosynthesis of SA and ROS, chloroplasts play a central role in plant immunity as integrators of environmental signals and transmitters of pro-defense signals (Serrano *et al.*, 2016; Lu & Yao, 2018). Stromules are stroma filled tubules that extend from chloroplasts and are induced in several different processes, including HR (Caplan *et al.*, 2015). Stromules are involved in retrograde signalling after pathogen invasion or light stress (Brunkard *et al.*, 2015; Caplan *et al.*, 2015) and in movement of chloroplasts within the cell (Kumar *et al.*, 2018).

It is established that SA is required for the restriction of pathogens during HR in various interactions, including plant-virus pathosystems (Mur *et al.*, 2008; Baebler *et al.*, 2014; Künstler *et al.*, 2016; Calil & Fontes, 2017). Several studies have also pointed to the essential role of apoplastic ROS leading to redox state changes in the cytoplasm in HR-conferred virus resistance (Hernández *et al.*, 2016; Lukan *et al.*, 2020). Although increases in chloroplastic ROS concentration have been observed in different incompatible plant-pathogen interactions (Zurbriggen *et al.*, 2010), the role of chloroplastic ROS during HR remains largely unknown. It has been suggested that chloroplastic ROS is involved in the signalling for and/or execution of HR cell death in incompatible interactions (Liu *et al.*, 2007; Zurbriggen *et al.*, 2009; Straus *et al.*, 2010; Ishiga *et al.*, 2011; Kim *et al.*, 2012; Xu *et al.*, 2019; Lukan *et al.*, 2020). Zurbriggen *et al.* (2009) suggested that ROS generated in chloroplasts during non-host interaction are essential for the progress of PCD, but do not contribute to the induction of pathogenesis-related genes or other signalling components of the response, including SA signalling. In contrast, Ochsenbein *et al.* (2006) found that chloroplastic singlet oxygen (^1^O_2_) can activate SA-mediated signalling, although SA is not required for ^1^O_2_-mediated cell death (Ochsenbein *et al.*, 2006). Straus *et al.* (2010) suggested that chloroplastic ROS acts as a flexible spatiotemporal integration point leading to opposite SA signalling reactions in infected and surrounding tissue (Straus *et al.*, 2010).

Moreover, recent evidence suggests that chloroplastic ROS might, in addition to signalling in HR cell death, also be involved in controlling plant immune responses by reprogramming transcription of genes involved in response to pathogen attack as one of the retrograde signals (Ochsenbein *et al.*, 2006; Lee *et al.*, 2007; Maruta *et al.*, 2012; Nomura *et al.*, 2012; Sewelam *et al.*, 2014; Pierella Karlusich *et al.*, 2017). For example, inducible silencing of chloroplastic ascorbate peroxidase increased H_2_O_2_ production in chloroplasts, which activated SA biosynthesis and SA-inducible gene expression (Maruta *et al.*, 2012).

To decipher the role of chloroplastic ROS in HR-conferred resistance, a detailed spatiotemporal analysis of chloroplast redox state in response to pathogen attack is needed. Nondestructive real-time measurement of redox state is feasible since Jiang *et al.* (2006) reported the use of redox state sensitive green fluorescent protein (roGFP). Measurement of roGFP fluorescence intensity following excitation at two different excitation maximum permits an evaluation of the relative proportion of roGFP in a reduced or oxidized state. By adding coding sequence for RuBisCO small subunit transit peptide to the roGFP coding sequence, pt-roGFP was constructed for measuring changes of redox state in chloroplasts (Stonebloom *et al.*, 2012). pt-roGFP also allows visualizing stromule formation and observation of redox state changes in stromules.

PVY, a member of the genus *Potyvirus*, is the most harmful virus of cultivated potatoes (Karasev & Gray, 2013) and is among top 10 most economically important plant viruses overall (Quenouille et al., 2013). In response to PVY, HR in potato cv. Rywal is manifested as the formation of necrotic lesions on inoculated leaves three days post inoculation and the virus is restricted to the site of inoculation (Szajko *et al.*, 2008). We have shown previously that SA regulates HR-conferred resistance in a spatiotemporal manner and proposed the role of chloroplastic ROS as a signal orchestrating PCD (Lukan *et al.*, 2020). However, the potential role of chloroplastic ROS in viral arrest has not yet been deciphered.

Here, we developed a protocol for confocal imaging complemented by custom image analysis scripts to simultaneously interrogate the chloroplast redox state and stromule formation in proximity to the cell death zone with spatiotemporal resolution. Using this protocol, we show that chloroplasts are strongly oxidized in the cells adjacent to cell death zone. More intriguingly, we detected individual cells with chloroplasts in moderately oxidized redox state close to the cell death zone as well as further away from it, indicating the role of chloroplastic ROS in signalling for HR-conferred resistance. As stromules are known to be involved in communication between chloroplast and nuclei, we have also assessed the frequency of stromule formation around the cell death zone in spatiotemporal manner. To decipher the link between SA signalling and chloroplast redox state and the link between SA signalling and stromule formation, we also analyzed the responses to PVY infection in SA-depleted transgenic counterpart (pt-roGFP-NahG). The presence of individual cells with moderately oxidized chloroplasts in the vicinity of the cell death zone or more distant to the cell death zone was sparse in SA-deficient plants, further supporting the hypothesis that these cells are involved in the signalling in HR-conferred resistance.

## Results

### Chloroplast redox state is highly oxidized in the cells surrounding the cell death zone

Our previous studies of HR in potato – PVY interaction suggested the role of chloroplastic ROS as a signal orchestrating PCD (Lukan *et al.*, 2020). To further explore the role of chloroplastic ROS in PCD induction in a spatiotemporal manner, we constructed transgenic plants of cv. Rywal with redox state sensitive GFP (roGFP) targeted to chloroplasts (hereafter pt-roGFP). Measurement of roGFP emission following excitation at 405 and 488 nm in constructed transgenic plants permits an evaluation of chloroplast relative redox state as the relative proportion of roGFP in a oxidized or reduced state (405/488 ratio) (Jiang *et al.*, 2006). Transgenic lines (pt-roGFP L2, L4 and L15) with strong and stable roGFP fluorescence in the chloroplasts were selected for further work (Supplemental Data Set 1).

To determine if chloroplastic ROS production is spatially and/or temporally regulated around the cell death zone, we measured the relative redox state in the chloroplasts of pt-roGFP plants in the areas adjacent to the lesion (ROI1) and adjacent to ROI1 (ROI2) at 3, 5 and 7 days post inoculation (dpi) with PVY, as well as in the area distant from the lesion (CTR) at 7 dpi (Figure 1a). Within the ROI1, chloroplasts in the cells surrounding the cell death zone were highly oxidized, in contrast to more distant cells (more distant from cell death zone within ROI1, in ROI2, and in CTR), where the chloroplastic redox state was more reduced (Figure 1b). As a control of normal physiological redox state in potato chloroplasts, we also measured chloroplast redox state in mock-inoculated pt-roGFP plants at 3 and 7 dpi (Figure 1a). The chloroplast redox state was highly uniform across all analysed mock samples and comparable to the redox state of chloroplasts in cells distant to cell death zone in PVY-infected plants (Figure 1b, right panel).

**Figure 1:**
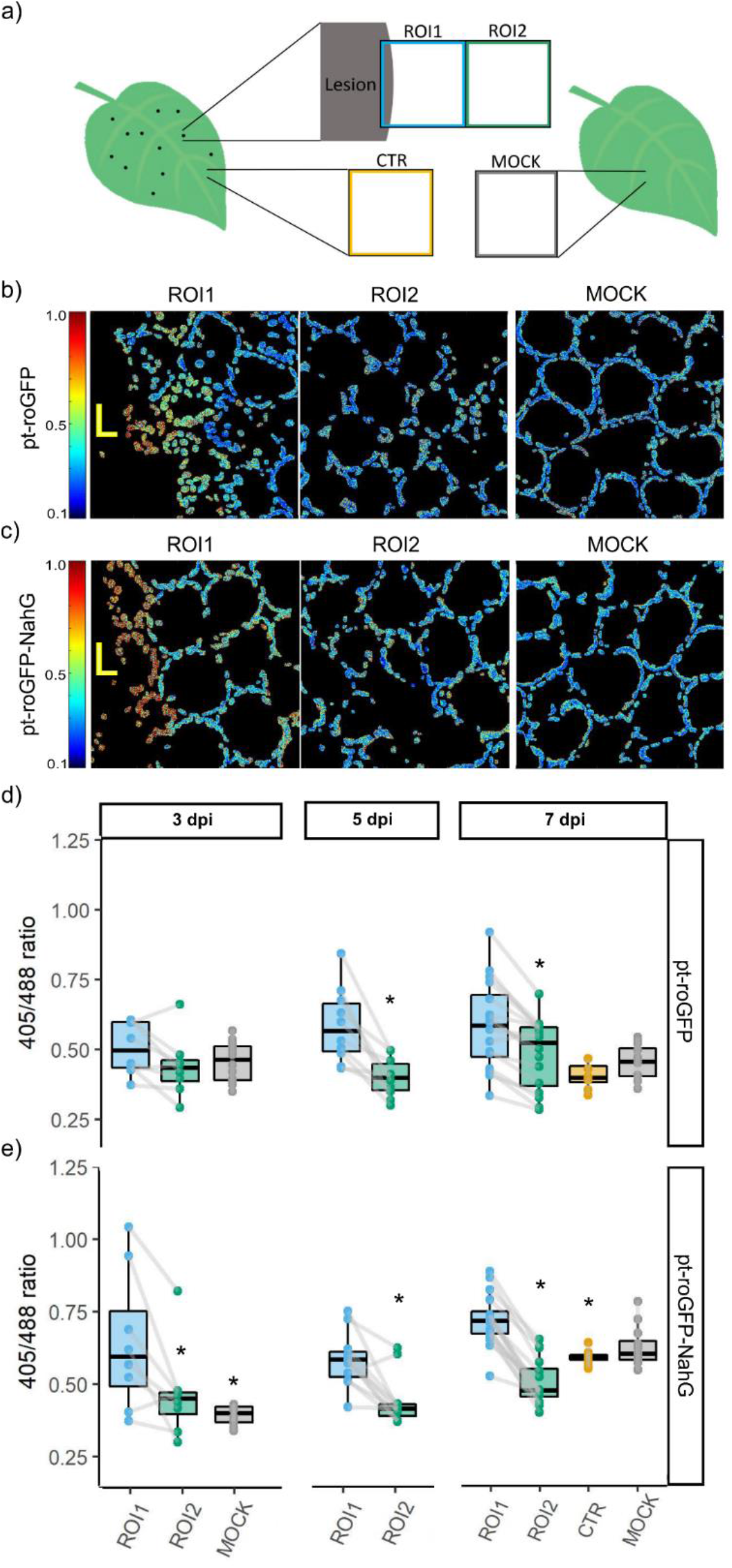
Chloroplast redox state is spatially regulated around the cell death zone during PVY-induced HR. a) After inoculating leaves with PVY, the chloroplast redox state was measured in transgenic sensor plants (pt-roGFP and SA-deficient pt-roGFP-NahG) with redox state sensitive GFP (roGFP) targeted to chloroplasts. Chloroplast redox state was measured in three regions: the area adjacent to the lesion (ROI1), the area adjacent to ROI1 (ROI2), and an area distant from the lesion (CTR) on the same leaf. As a negative control, chloroplast redox state was also measured in mock-inoculated plants. b, c) Visual presentation of the ratios of fluorescence intensities of roGFP after excitation with 405 and 488 nm laser (405/488 ratio, chloroplast redox state) presented on the rainbow scale for pt-roGFP L2 (b) and pt-roGFP-NahG L2 (c) at 5 days post inoculation in ROI1, ROI2 and MOCK (from left to right). Higher ratios denote chloroplasts in a more oxidized state (red). Chloroplasts in the cells surrounding the cell death zone are highly oxidized, in contrast with more distant cells. d, e) Chloroplast redox state in the above-mentioned leaf areas in pt-roGFP L2 (d) and pt-roGFP-NahG L2 (e) plants 3, 5, or 7 days post-inoculation. Results are presented as boxplots with 405/488 ratios of each measured ROI shown as dots (Exp3NahG and Exp5NT in the Supplemental Data Set 1). Grey lines connect ROI1 and ROI2 pairs for each lesion. Asterisks denotes statistically significant difference between the marked regions (ROI2, CTR or MOCK) and ROI1.

Using custom image analysis scripts for quantifying chloroplast redox state in the regions surrounding the cell death zone with spatiotemporal resolution and statistics, we show that chloroplast redox state is in ROI1 statistically significantly more oxidized from the chloroplast redox state in ROI2 and CTR (Figure 1d) at late time points after inoculation (5 and 7 dpi). The experiment was performed three times independently on three transgenic lines (Redox Exp6NT in Figure Supplemental Figure 1b, Supplemenal Data Set 1, Supplemental Data Set 2).

### Individual cells with moderately oxidized chloroplasts spread farther away from the cell death zone

On the edge of the cell death zone, within the ROI1, the majority of chloroplasts in strongly oxidized redox state were disordered (Figure 2; marked with L), most probably belonging to the mesophyll cells in transition between normal cells and cells in PCD as determined by electron microscopy in our previous work (Lukan et al., 2020). However, we also observed chloroplasts in moderately oxidized redox state that were well arranged, following the shape of a normal mesophyll cell (Figure 1b). Moreover, we often observed such individual cells, with chloroplasts in moderately oxidized redox state, farther from the cell death zone in ROI2 (Figure 2, Supplemental Figure 2). On the other hand, no such cells were observed in mock-inoculated plants (Figure 1b).

**Figure 2:**
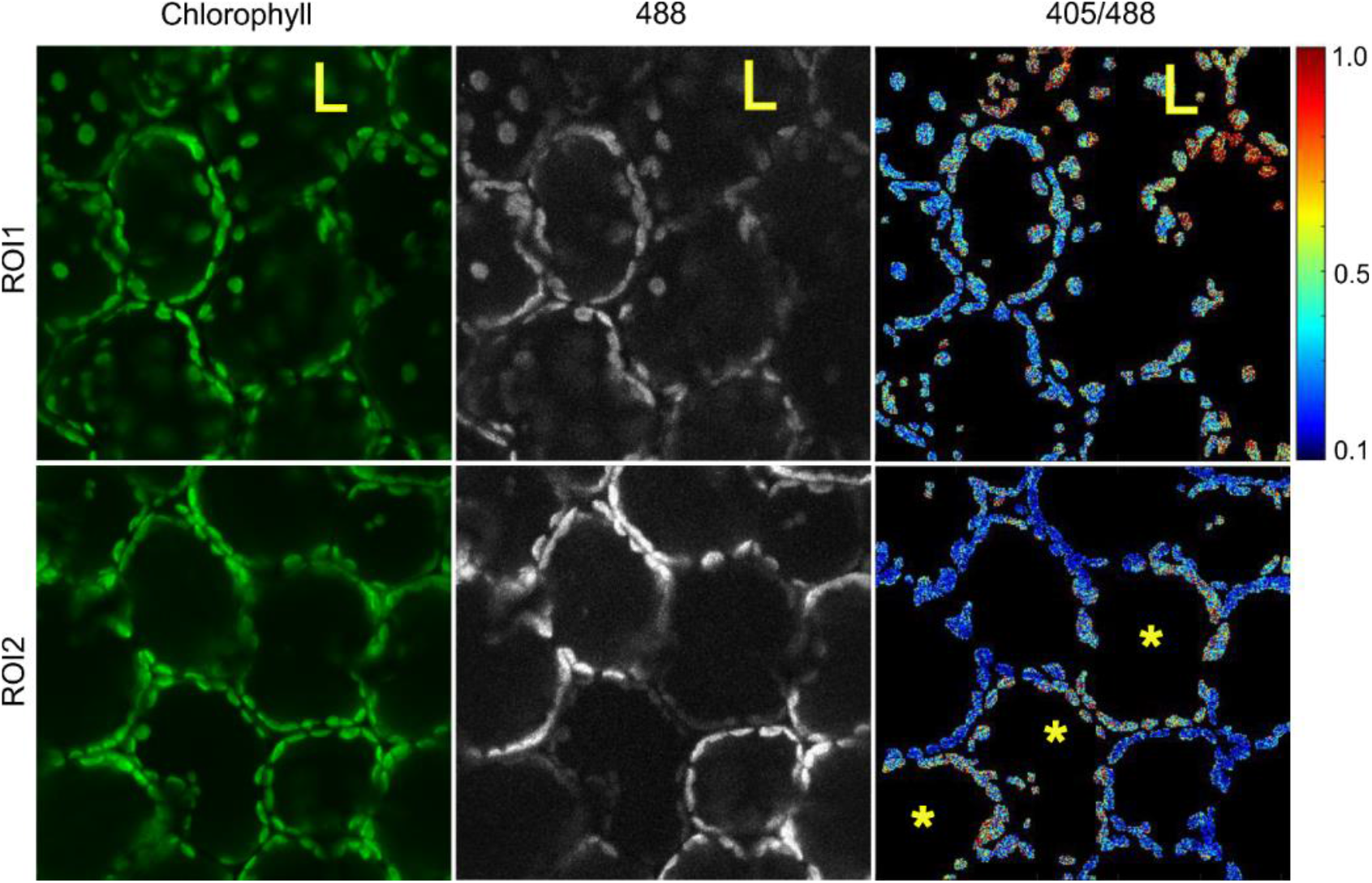
Differential chloroplast redox state around the HR cell death zone. Beside strongly oxidized chloroplasts (red) in the proximity of cell death zone (L), individual cells with moderately oxidized chloroplasts (asterisks) can also be observed farther from the cell death zone (ROI2). Upper and lower panel present two consecutive regions in cell death zone proximity (ROI1, ROI2). All images were taken in pt-roGFP plants at 5 days post inoculation. Cells with oxidized chloroplasts farther away from the cell death zone are marked with asterisks. Cell death zone is marked with L. ROI1: region of interest adjacent to cell death zone, ROI2: region of interest adjacent to ROI1 (see Figure 1a). Chlorophyll: chlorophyll fluorescence, 488: roGFP fluorescence after excitation with 488 nm laser line showing reduced roGFP (brighter chloroplasts are more reduced), 405/488: relative redox state (405/488 ratio) presented on a rainbow scale. Higher ratios denote more oxidized chloroplasts (red).

The measured variability of chloroplast redox state was in ROI1 and ROI2 higher than in mock samples (Figure 1d), which could well be due to described specific features of HR response, namely the highly oxidized chloroplasts in cells of PCD transition zone and the occurrence of cells with moderately oxidized chloroplasts farther away (Figure 1d, Figure 2). To confirm our hypothesis, we separated ROI1 and ROI2 into five regions (Bins), according to the distance from the cell death zone (Figure 3a). The results confirm our hypothesis as the chloroplast redox state is the most oxidized close to the cell death zone (Bin1) and is getting less oxidized with increasing distance from the cell death zone at different time points (Figure 3b and d, Supplemental Figure 3). In the most distant Bin from the cell death zone (Bin5 of ROI1), the redox state stabilizes and remains comparable in all five Bins of ROI2 (Figure 3b and d). The moderate variability within these Bins can be attributed to moderately oxidized chloroplasts belonging to individual cells observed farther away from the cell death zone (Figure 2a, Figure 3d).

**Figure 3:**
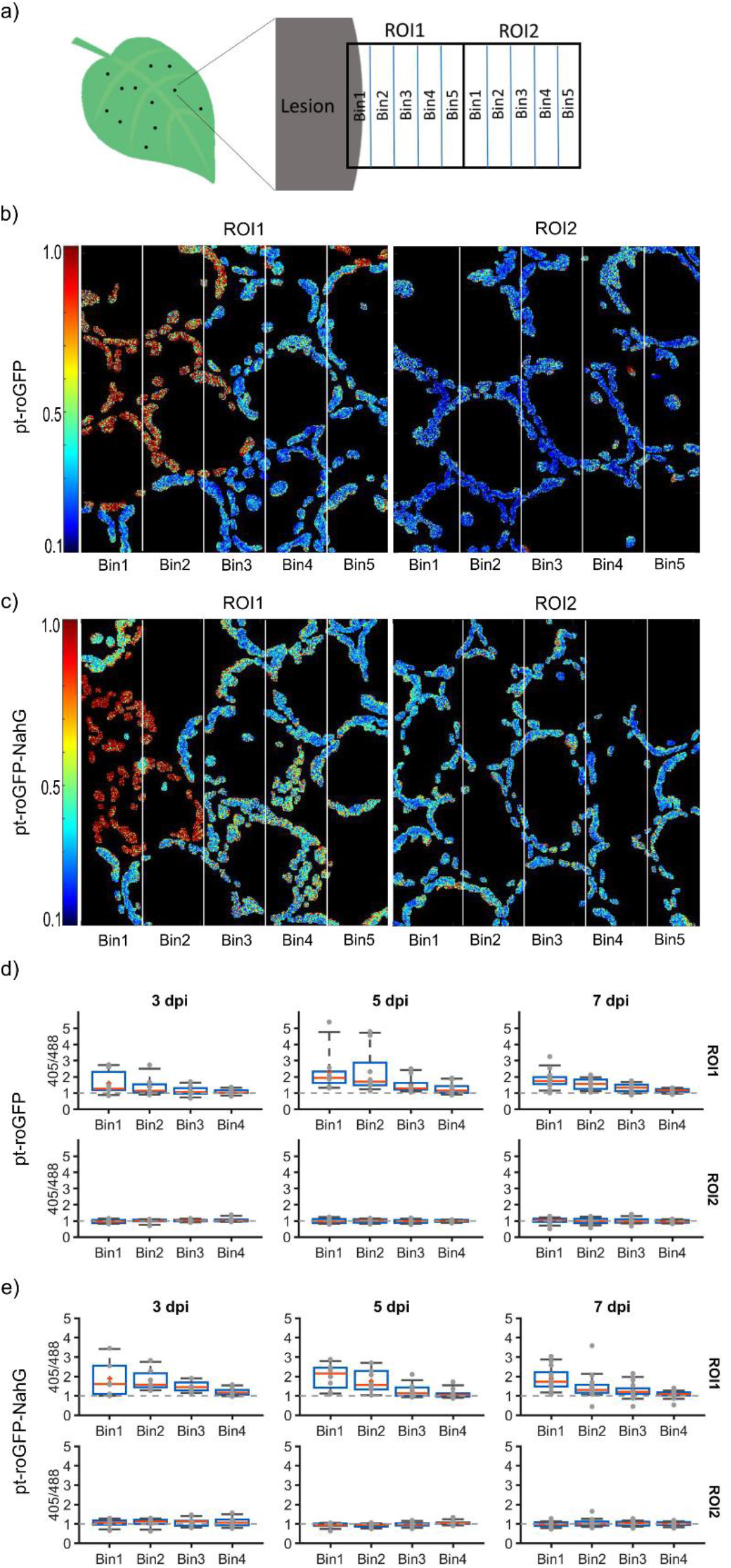
Chloroplast redox state is more oxidized around the cell death zone and is getting more reduced with increasing distance from the cell death zone. a) Chloroplast redox state, measured in redox state sensor plants (pt-roGFP and SA-deficient pt-roGFP-NahG) with redox state sensitive GFP (roGFP) targeted to chloroplasts following PVY inoculation in the area adjacent to the lesion (ROI1) and adjacent to ROI1 (ROI2) were separated into five regions (Bins) according to the distance from the cell death zone. b, c) Visual presentation of the ratios of fluorescence intensities of roGFP after excitation with 405 and 488 nm laser (405/488 ratio, chloroplast redox state) presented on the rainbow scale for pt-roGFP L2 (b) and pt-roGFP-NahG L2 (c) at 5 days post inoculation. Higher ratios denote more oxidized chloroplasts (red). d, e) The normalized 405/488 ratio in five Bins of ROI1 (top panel) and ROI2 (bottom panel) in pt-roGFP L2 (d) and pt-roGFP-NahG L2 (e). The 405/488 ratios in each of the first four Bins were normalized to the 405/488 ratio from Bin 5, which was set to 1 (dotted line) in both ROIs. 405/488 ratio in Bin 5 of ROI1 was similar to 405/488 ratio in Bin 5 of ROI2, allowing for comparisons within and between ROIs. Results are presented as boxplots, with normalized 405/488 ratio for each Bin shown as dots (Redox Exp3NahG and Exp5NT in the Supplemental Data Set 1 and Supplemental Data Set 3). L – transgenic line.

### Presence of cells with oxidized chloroplasts farther away from the cell death zone suggest their role in signal transmission in HR-conferred resistance

In order to study the link between SA signalling and chloroplast redox state in PCD and resistance, we introduced roGFP also in SA-depleted transgenic counterpart NahG-Rywal (hereafter pt-roGFP-NahG). Transgenic lines (pt-roGFP-NahG L2 and L7) with strong and stable GFP fluorescence in the chloroplasts were selected for further work. We next followed chloroplast redox state after viral infection in different regions, close or farther away from the cell death zone at three time points. Similarly as in pt-roGFP plants (Figure 1b), within the ROI1, chloroplasts in the cells surrounding the cell death zone were highly oxidized, in contrast to more distant cells (more distant from cell death zone within ROI1 and in ROI2), where redox state was reduced (Figure 1c). This was further confirmed by image analysis, as chloroplast redox state was more oxidized in ROI1 and was statistically significantly different from the chloroplast redox state in ROI2 at all analysed time points (Figure 1e, Supplemental Figure 1a, Supplemental Data Set 1, Supplemental Data Set 2). We conclude, that chloroplast redox state is precisely spatially regulated around the cell death zone both in pt-roGFP and SA-deficient pt-roGFP-NahG plants, as it is more oxidized adjacent to the cell death zone and is getting more reduced with increasing distance from the cell death zone in both genotypes. Interestingly however, the cells with moderately oxidized chloroplasts, farther away from the lesion, were rarely observed in SA-deficient plants. We thus hypothesize on their signalling role in HR-conferred resistance.

### Spatiotemporal regulation of stromule formation around the cell death zone is SA-signalling-dependent

The individual cells with oxidized chloroplasts farther away from the cell death zone indicate on their signalling role in HR-conferred resistance. This hypothesis is further supported by the presence of chloroplasts with stromules in the proximity of cells with oxidized chloroplasts (Figure 4a and b, Supplemental Figure 4). Interestingly, chloroplasts with stromules were in a strongly reduced state (Figure 4a and b, Supplemental Figure 4).

**Figure 4:**
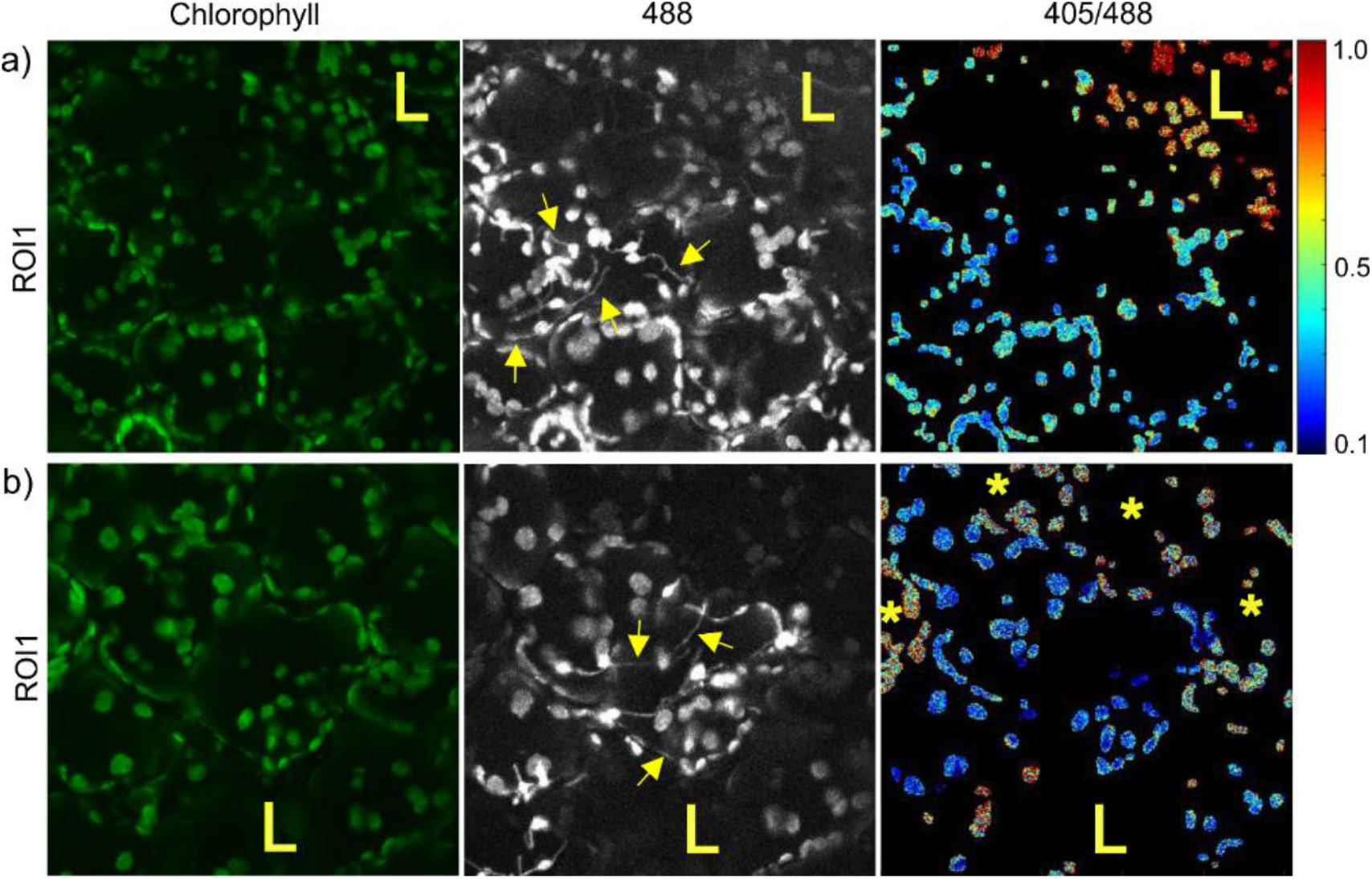
Stromule occurrence around the HR cell death zone. a, b) Chloroplasts with stromules (arrows) in the proximity of cells with oxidized chloroplasts on the edge of cell death zone (a) or farther away from the cell death zone (b). Chloroplasts with stromules were in a strongly reduced redox state (high signal in 488 channel, thus not visible on 405/488 ratio images) and were often in close proximity to the cells with oxidized chloroplasts. All images were taken in pt-roGFP plants at 5 days post inoculation. Stromules are marked with arrows. Cells with oxidized chloroplasts farther away from the cell death zone are marked with asterisks. Cell death zone is marked with L. ROI1: region of interest adjacent to cell death zone (see Figure 1a). Chlorophyll: chlorophyll fluorescence, 488: roGFP fluorescence after excitation with 488 nm laser line showing reduced roGFP (brighter chloroplasts are more reduced), 405/488: relative redox state (405/488 ratio) presented on a rainbow scale. Higher ratios denote more oxidized chloroplasts (red).

To further investigate the potential role of stromules in HR-conferred resistance, we performed experiments in which we followed stromule formation in the areas adjacent to the lesion (ROI1) and adjacent to ROI1 (ROI2) (Figure 5a) at three time points following inoculation (3, 4 and 5 dpi). We observed increased stromule formation frequency after inoculation at all time points, which was even more pronounced in the cell death zone proximity in pt-roGFP plants (Figure 5b, c and d). When comparing the frequency of stromules between ROI1 and ROI2, in pt-roGFP plants, the difference was statistically significant at 4 and 5 dpi. On the other hand, in pt-roGFP-NahG plants, the number of stromules was similar between ROI1 and ROI2 at all time points (Figure 5d, Supplemental Data Set 4, Supplemental Data Set 5). The frequency of stromules of PVY-inoculated plants was higher if compared to random regions on the leaf of mock-inoculated plants at all time points in pt-roGFP plants (Figure 5d, Supplemental Data Set 4, Supplemental Data Set 5). The induction of stromules was however less pronounced in pt-roGFP-NahG plants. We thereby conclude that in PVY-induced HR in potato, stromule formation is spatiotemporaly regulated around the cell death zone and is SA signalling-dependent.

**Figure 5:**
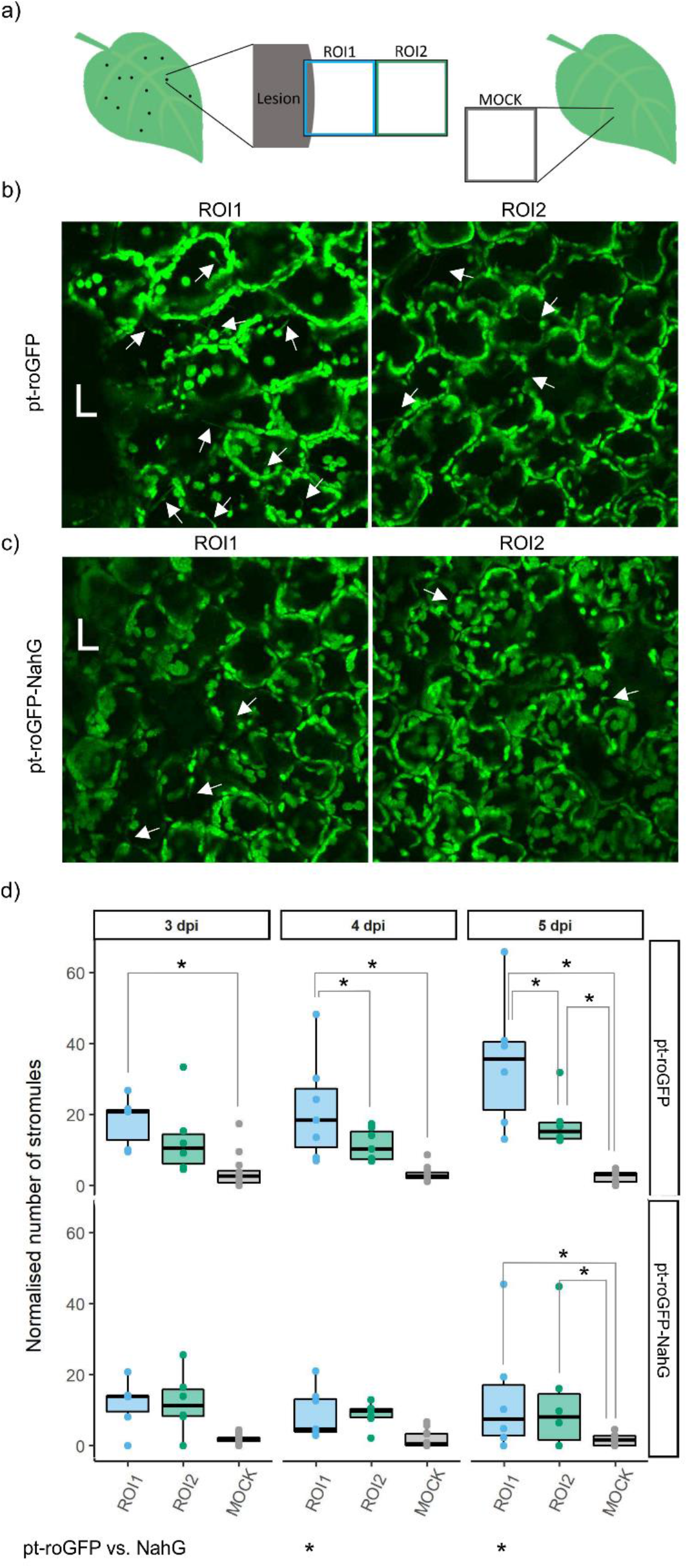
Spatiotemporal regulation of stromule formation around the cell death zone is SA-signalling-dependent. a) Stromule formation was followed in redox state sensor plants (pt-roGFP and SA-deficient pt-roGFP-NahG) with redox state sensitive GFP targeted to chloroplasts following PVY inoculation in the area adjacent to the lesion (ROI1) and adjacent to ROI1 (ROI2). As a control, stromule formation was also followed in mock-inoculated plants (MOCK). b, c) Confocal image showing the difference in the number of stromules between ROI1 (left) and ROI2 (right) in pt-roGFP (b) and pt-roGFP-NahG (c). After inoculation, we observed stromule formation, which was even more pronounced in the cell death zone proximity in pt-roGFP plants. Arrows show stromules. d) Normalised number of stromules was calculated by dividing the number of stromules by the number of chloroplasts counted in the above mentioned leaf areas in pt-roGFP L2 and pt-roGFP-NahG L2 transgenic lines at 3, 4 and 5 days post inoculation (Exp6NTNahG in the Supplemental Data Set 4). Results are presented as boxplots with normalised numbers of stromules for each ROI shown as dots (Stromules Exp6NTNahG in the Supplemental Data Set 4). Asterisks denotes statistically significant differences between regions (shown on the boxplots connecting lines; see Stromules Exp6NTNahG in the Supplemental Data Set 5 for p values) or between genotypes (pt-roGFP vs. NahG, shown for each region for each dpi; see Stromules Exp6NTNahG in the Supplemental Data Set 5 for p values). The difference in the number of stromules between ROI1 and ROI2 was in pt-roGFP statistically significant at 4 and 5 dpi, while in pt-roGFP-NahG, the number of stromules was similar between ROI1 and ROI2 at all dpi.

### Stromules are induced in close proximity to virus multiplication zone

As in ROI2 the frequency of stromules was still higher than in mock samples, we next followed stromule formation in four consecutive regions adjacent to the cell death zone at different time points following inoculation in both genotypes (ROI1-ROI4; Supplemental Figure 5a, b, d). We confirmed the results from the previous experiment showing that stromule formation is differently spatiotemporally regulated between genotypes. In pt-roGFP-NahG, the difference in stromule frequency between regions was observed only later after inoculation (5-6 dpi) and farther from the cell death zone, while in pt-roGFP, the difference was visible already at early time points (4 and 5 dpi) and closer to the cell death zone (Figure 5, Supplemental Figure 5a, b, d, Supplemental Data Set 4, Supplemental Data Set 5).

In our previous study, we observed PVY-GFP accumulation in the cells on the edge of the cell death zone at all analyzed time points, most frequently at 4 dpi, in non-trangenic plants, but the virus did not spread farther. In NahG plants, we observed concentric virus spread to adjacent cells, at 6 and 7 dpi the virus reached outer borders of ROI2 (Lukan *et al.*, 2018). This pattern correlates with stromules frequency in cell death zone proximity, which is in pt-roGFP statistically higher in ROI1, while in pt-roGFP-NahG, both ROI1 and ROI2 have higher stromule frequency compared to ROI3, ROI4 or MOCK (Figure 5d, Supplemental Figure 5a, b, d). Stromules frequency was not statistically significantly different between ROI1 and ROI2 in pt-roGFP at 3 dpi most likely as PVY-GFP is not accumulated to the extent that could be detected in the cells adjacent to the cell death zone at that time point (Lukan *et al.*, 2018).

To determine the relation between chloroplast redox state, stromule frequency and virus multiplication, we followed the cells on the edge of the virus multiplication zone in pt-roGFP at 4 dpi and in pt-roGFP-NahG at 7 dpi (Figure 6a and c, Supplemental Data Set 1). In pt-roGFP, in the cell next to the cell in which we observed PVY-GFP accumulation, we observed reduced chloroplasts with stromules (Figure 6b). On the other hand, as expected, we observed oxidized chloroplasts on the edge of the cell death zone (Figure 6b). Similarly, in pt-roGFP-NahG, we observed cells with reduced chloroplasts with stromules, in the proximity of the cell in which we observed PVY-GFP accumulation (Figure 6d). However, in neither of the studied genotypes we observed any cells with moderately oxidized chloroplasts close to the edge of virus multiplication. We conclude that stromules formation is also a response to virus multiplication, while there is no direct connection between virus multiplication and the cells with moderately oxidized chloroplasts.

**Figure 6:**
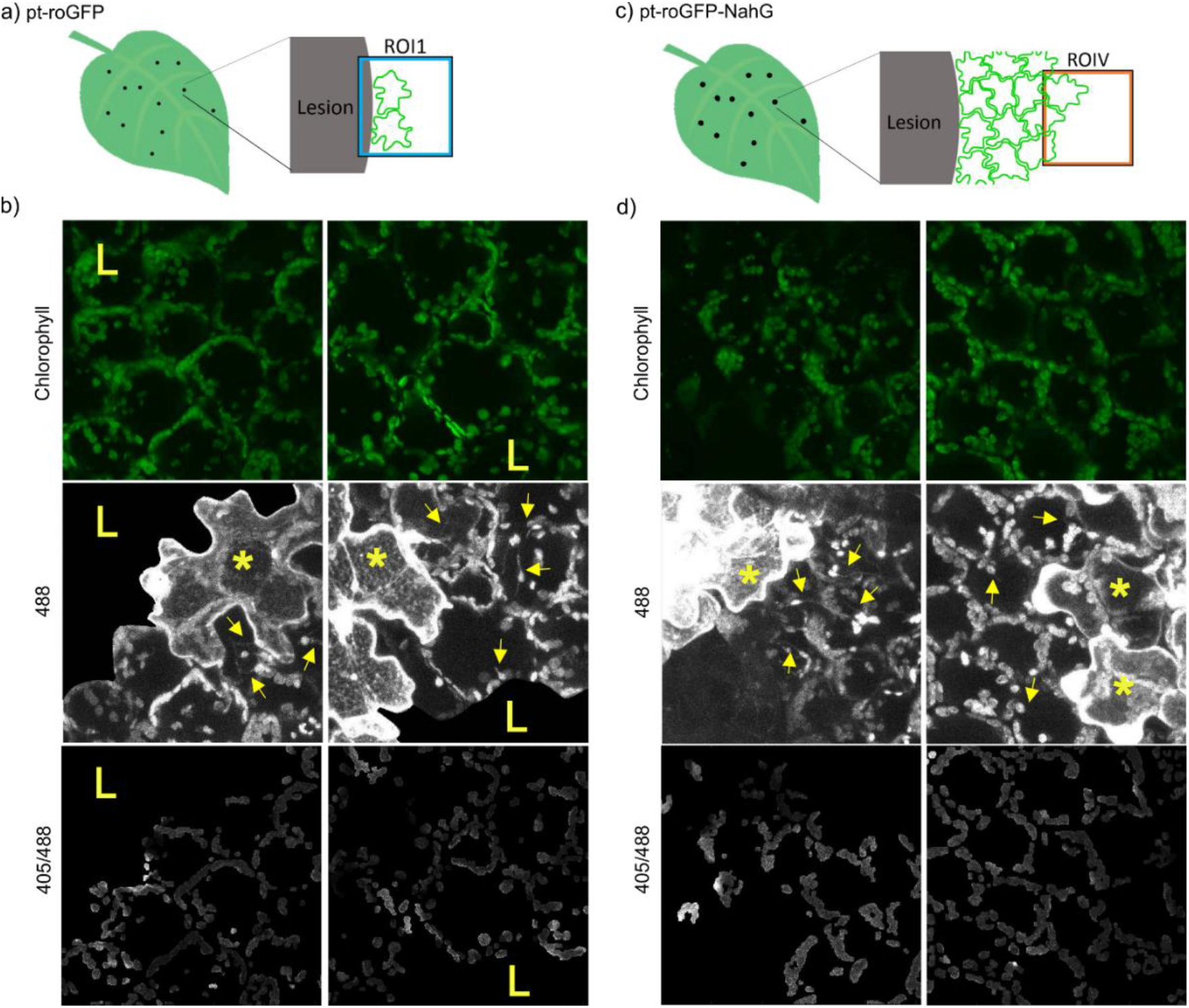
Stromules are induced in close proximity to the virus multiplication zone. a, c) Stromule formation and relative chloroplast redox state were followed in pt-roGFP on the edge of cell death zone (ROI1) in the cells adjacent to the cell in which we observed PVY-GFP accumulation at 4 dpi (a) or in pt-roGFP-NahG on the edge of the virus multiplication zone (ROIV) in the cells adjacent to the cell in which we observed PVY-GFP accumulation at 7 dpi (c). b and d) 488: stromules (arrows) are induced adjacent to the cell in which we observed PVY-GFP accumulation (asterisks), 405/488: chloroplasts are oxidized on the edge of cell death zone (L), but not in the cells adjacent to the cells with PVY-GFP accumulation (asterisks). Due to high background signal in the cell death zone, as a result of virus derived GFP fluorescence, GFP signal or 405/488 ratios are not shown in the cell death zone in images 488 and 405/488, respectively. Left and right panels present different ROIs. Stromules are marked with arrows. Cells in which we observed PVY-GFP accumulation are marked with asterisks. Cell death zone is marked with L. ROI1: region of interest adjacent to cell death zone, ROIV: region of interest adjacent to the virus multiplication zone. Chlorophyll: chlorophyll fluorescence, 488: roGFP fluorescence after excitation with 488 nm laser showing reduced roGFP and GFP fluorescence as a result of PVY-GFP multiplication (asterisks), 405/488: relative redox state (405/488 ratio) presented on a greyscale (higher ratios denote more oxidized chloroplasts (bright)).

## Discussion

PCD has been associated with increased chloroplastic ROS production (Liu et al., 2007; Straus et al., 2010; Wagner et al., 2004). Recently, however, generation of chloroplastic ROS was proposed as a signal orchestrating PCD, rather than being only the consequence of PCD (Zurbriggen et al., 2010; Karlusich et al., 2017; Su et al., 2018). Our results, showing that the disordered chloroplasts were oxidized in the cells adjacent to the cell death zone (Figure 3b and c top panel, d and e left image, Supplemental Figure 3) are in agreement with the results of above-mentioned studies suggesting the involvement of chloroplastic ROS as a signal orchestrating PCD or being the consequence of PCD. In addition, close to the edge of the cell death zone (ROI1, Figure 3d, e), as well as farther from the cell death zone (ROI2, Figure 2a, Supplemental Figure 2, Supplemental Data Set 1), we observed individual mesophyll cells with moderately oxidized chloroplasts. These cells were sparse in SA-deficient plants, which indicates on their role in signal transmission in HR-conferred resistance. They could, therefore, be called “signalling cells”. This is further supported by the results of several studies suggesting that decreased chloroplastic ROS production leads to compromised resistance (Su *et al.*, 2018; Xu *et al.*, 2019; Schmidt *et al.*, 2019).

Chloroplast redox state has previously been studied in HR-PCD triggered by TMV effector p50 using HyPer2 H_2_O_2_ sensor (Caplan *et al.*, 2015). They detected higher levels of ROS in chloroplasts at 28 h post induction of p50 accumulation. However, using such experimental approach, no information on spatial distribution of responses could be obtained. On the other hand, spatiotemporal studies were performed to follow chloroplast redox state in relation to light stress (Stonebloom *et al.*, 2012; Brunkard *et al.*, 2015; Exposito-Rodriguez *et al.*, 2017). Our study thus brings new perspective into signalling in HR-resistance.

We show that more oxidized chloroplast redox state is correlated with a higher number of stromules, as in pt-roGFP plants more stromules were detected in ROI1 with more oxidized chloroplast redox state, compared to ROI2 (Figure 5). These results are in agreement with the results of several studies showing that treatment with exogenous H_2_O_2_, induction of chloroplastic ROS by light or NADPH-dependent thioredoxin reductase silencing resulted in stromules induction (Brunkard *et al.*, 2015; Caplan *et al.*, 2015; Barton *et al.*, 2018; Ding *et al.*, 2019). Interestingly, however, our results show that the cells with the highest frequency of stromules are in close proximity of so-called signalling cells (cells with moderately oxidized chloroplasts farther from the cell death zone) and their chloroplasts are reduced (Figure 2c). This is in disagreement with the results of above mentioned studies (Brunkard *et al.*, 2015; Caplan *et al.*, 2015; Barton *et al.*, 2018; Ding *et al.*, 2019). It is possible, however, that our findings are specific for cell-to-cell signalling in plant-virus interaction, which was so far not studied in this aspect.

Stonebloom *et al.* (2012) suggested that ROS production in plant organelles triggers intracellular communication pathways that alter intercellular communication via plasmodesmata (PD). They showed that oxidized plastids resulted in decreased PD transport, while reduced plastids facilitate PD transport. Thus, we could assume that the highly oxidized chloroplasts at the edge of cell death zone have PD closed to prevent the spread of the pathogen. Interestingly, the cells with high frequency of stromules that we observed in cell death zone proximity or in the proximity of the cell in which we observed PVY-GFP accumulation, had reduced chloroplasts (Figure 6), meaning they could have increased PD transport. This is in accordance with Caplan *et al.* (2015), who suggested that stromules function is HR-PCD signalling. Their experimental approach with transient expression of effector proteins, however, does not enable distinction between signalling for PCD and signalling for viral arrest. Stromules were shown to contribute to immune response also in susceptible *N. benthamiana* - *Phytophthora* interaction (Toufexi et al., 2019).

Maruta *et al.* 2012 suggested that the chloroplastic H_2_O_2_ and SA activate each other and that this positive feedback loop is involved in the plant response to biotic stress. In our study, no major differences were observed between the chloroplast redox state of SA-deficient pt-roGFP-NahG compared to pt-roGFP plants (Figures 1 and 3), except for the frequency of previously mentioned signalling cells with moderately oxidized chloroplasts (Figure 2). Additionally, we studied SA involvement in stromule formation. The number of stromules in ROI1 was statistically significantly higher in pt-roGFP if compared to SA-deficient pt-roGFP-NahG plants (Figure 5). This is in agreement with results from Caplan *et al.* (2015), who showed that exogenous application of SA analog results in stromules induction when HR is triggered by pathogen effectors. However, since in our pathosystem the number of stromules was in PVY-inoculated SA-deficient plants higher if compared to mock-inoculated plants, but lower if compared to PVY-inoculated pt-roGFP plants (Figure 5c), we conclude that stromules induction does not depend solely on SA. The increase in stromule formation observed in SA-deficient plants in ROI2 (Supplemental Figure 5) was however linked to the border of virus multiplication zone (Figure 6). The only viral amplification occurs in the cells adjacent to cell death zone in pt-roGFP plants (Lukan, 2018), corresponding to induction of stromules in ROI1. Therefore, we conclude that the stromules are in our pathosystem involved in signalling on the front of virus multiplication and that this response is SA-signalling dependent (Figure 4).

Our results show that HR-conferred resistance to the virus is mediated by so-called signalling cells, cells with moderately oxidized chloroplasts farther away from the cell death zone, that evidence resistance signal transmission to neighboring tissue. Induction of stromule formation in the vicinity of those cells further supports their role in HR-conferred resistance. However, as some pathogens target chloroplast-related functions to subvert host defences and promote virulence (Serrano *et al.*, 2016), it is plausible that the redox state changes that we detected are not solely the results of plant response to pathogen attack, but rather the results of two intertwined processes – plant response and virus manipulation of plant immune response.

## Methods

### Generation of redox state sensor plants

Potato (*Solanum tuberosum* L.) transgenic lines pt-roGFP and pt-roGFP-NahG were prepared by stable transformation of resistant cv. Rywal and susceptible NahG-Rywal (the depletion of SA renders NahG plants susceptible (catechol independent); Baebler *et al.*, 2014) with a construct pCAMBIA1304_pt-roGFP2 (Stonebloom *et al.*, 2012). pCAMBIA1304_pt-roGFP2 was electroporated into *Agrobacterium tumefaciens* LBA4404 (Eppendorf Electroporator 2510) following manufacturer’s protocol at 2000 V. The transformed bacteria were used for the transformation of cv. Rywal and NahG-Rywal stem internodes from *in vitro* plantlets as described elsewhere (Baebler *et al.*, 2014), with some modifications (Supplemental Method 1). Well-rooted hygromycin-resistant plants were sub-cultured to produce plantlets of the independently transformed lines. The selection of transgenic lines with strong and stable GFP fluorescence in chloroplasts was performed by confocal microscopy (see below). To confirm that the detected signal is specific for GFP emission, we used cv. Rywal and NahG-Rywal plants as a control.

Transgenic lines were grown in stem node tissue culture. Two weeks after node segmentation, they were transferred to soil in a growth chamber and kept under controlled environmental conditions as described elsewhere (Baebler *et al.*, 2009).

### Construction of PVY-N605(123)-GFP infectious clone

PVY-N605(123)-GFP was constructed by inserting GFP coding sequence between the coding sequences for viral proteins NIb and CP in PVY-N605(123) infectious clone (Bukovinszki *et al.*, 2007) using a similar design as Rupar *et al.* 2015, allowing the GFP reporter to be excised from the polyprotein following translation.

GFP coding sequence was amplified from plasmid p2GWF7 (Karimi *et al.*, 2002) using GFPF40 and GFPR40 primers (see Supplemental Method 2) with Phusion polymerase (New England BioLabs) according to the provider’s instructions. The primers were designed with overhangs enabling addition of PVY-N605(123) annealing sequence and protease recognition site to the GFP sequence, making the amplicon serve as a megaprimer for restriction-free insertion with mutagenesis. Reaction was designed using the RF-Cloning online tool (Bond & Naus, 2012) and carried out using Phusion polymerase with 1:20 molar ratio of PVY-N605(123) plasmid to GFP amplicon (megaprimer). After *Dpn*I digestion, mutagenesis mixture was transformed into *E. coli* XL10-Gold Ultracompetent Cells following manufacturer’s instructions (Agilent). The transformants were plated on LB-agar with ampicillin selection and grown overnight at 37°C. Grown colonies were screened with colony PCR using CP-F and UnivR primers and KAPA2G Robust HotStart Kit (Agilent, Supplemental Method 2). Sanger sequencing of the selected clone confirmed correct sequence of the PVY-coding part and correct in-frame insertion of GFP coding sequence. Constructed PVY-N605(123)-GFP was coated onto gold microcarriers and used for *Nicotiana clevelandii* bombardment with Helios® gene gun (BioRad) according to Stare et al. (2020). Detailed PCR reaction conditions and cloning steps are given in Supplemental Method 2.

### Virus inoculation

Two to four weeks old potato plants in soil were inoculated with PVY^N-Wilga^ (PVY^N–Wi^; EF558545), PVY N605-GFP (Rupar *et al.*, 2015), PVY-N605(123)-GFP or MOCK, as described in Baebler *et al.*, 2009.

### Treatments with oxidant and reductant

A second and third leaf of two to four weeks old pt-roGFP and pt-roGFP-NahG potato transgenic lines were treated with an oxidant (200 mM solution of hydrogen peroxide (H_2_O_2_)), a reductant (200 mM solution of dithiothreitol (DTT)) or a control (bidistilled water (ddH_2_O)) by vacuum infiltration using a syringe. We followed relative redox state in the chloroplasts 30 minutes after the treatment, to confirm that redox state sensor roGFP is functional in the selected transgenic lines (Supplemental Figure 1, Supplemental Figure 6, Supplemental Data Set 1, and Supplemental Data Set 2).

### Confocal microscopy

Confocal microscope (Leica TCS LSI macroscope with Plan APO 5 x and 20 x objective, Leica Microsystems) was used to detect emission of chlorophyll or redox state sensitive GFP (roGFP) for the selection of transgenic plants, stromule counting and redox state detection.

To select potato transgenic lines with strong and stable roGFP fluorescence (emission window 505 – 520 nm) in the chloroplasts, plants from tissue cultures were analysed with 20 x objective, zoom factor set to 3.00, frame average to 2 and z-stack size adjusted to 10 steps to cover at least 30 μm of the mesophyll after excitation with 488 nm laser.

For redox state detection experiments, the emission of roGFP was followed after excitation with 405 nm (detection of oxidized roGFP) and 488 nm (detection of reduced roGFP) laser in the window between 505 and 520 nm. For stromules counting experiments, the emission of roGFP was measured after excitation with 488 nm laser line in the window between 505 and 530 nm. The background chloroplast fluorescence was excited with the 488 nm laser line and the emission was measured in the window between 690 and 750 nm. The fluorescence was followed on the upper side of the leaf which was detached from the plant on the particular day post inoculation (dpi) or 30 minutes after the treatment with an oxidant or a reductant. Regions of interest (ROI) were scanned unidirectionally with scan speed of 400 Hz. For the redox state detection experiments, fluorescence emissions were collected sequentially through three channels (roGFP fluorescence after excitation with 405 nm laser line, roGFP fluorescence after excitation with 488 nm laser line and chlorophyll fluorescence), while for stromules counting experiments through two channels (roGFP fluorescence after excitation with 488 nm laser line and chlorophyll fluorescence). ROI sizes and magnifications for individual experiments are specified in Supplemental Data Set 1 and Supplemental Data Set 4. The images were processed using Leica LAS X software (Leica Microsystems) to obtain maximum projections from Z-stacks for each of two or three channels.

In redox state detection experiments (Supplemental Data Set 1), PVY^N-Wilga^- or PVY-N605(123)-GFP-inoculated, MOCK-inoculated and oxidant-/reductant-treated plants were imaged with 20 x objective, zoom factor set to 3.00, frame average to 2. On MOCK-inoculated and oxidant-/reductant-treated plants ROIs were selected randomly on the leaf. On PVY^N-Wilga^-inoculated plants, two consecutive ROIs were imaged adjacent to the lesion (ROI1, ROI2) and one ROI was imaged distant from the lesion (CTR) (see Figure 1a). See Supplemental Data Set 1 for the number of analyzed plants, leaves, lesions and ROIs for each experiment for each treatment and time point. Z-stack size was adjusted to 10 steps to cover at least 30 μm of the mesophyll. On PVY-N605(123)-GFP -inoculated plants ROI1 was imaged adjacent to the lesion (ROI1) or on the front of the virus spread (ROIV) (see Figure 6 and Supplemental Data Set 1). For virus localization z-stack size was adjusted to 10 steps to cover the whole epidermis (~30 μm) and at least 30 μm of the mesophyll, while for redox state detection z-stack size was adjusted to 10 steps to cover at least 30 μm of the mesophyll (see comments in Supplemental Data Set 1). Due to high signal in 405 channel in the cell death zone and high signal in 488 channel in the chloroplasts in the cell in which we observed PVY-N605(123)-GFP accumulation, we only focused on the redox state and stromules formation in the cells adjacent to the cells in which we observed PVY-N605(123)-GFP accumulation.

In stromules counting experiments, PVY N605-GFP-inoculated plants were imaged. In Exp3NT and Exp5NahG (Supplemental Data Set 4), plants were imaged with 5 x objective, frame average was set to 3 and z-stack was adjusted to 12-15 steps (1.50 - 2.00 μm per step). Four consecutive ROIs were imaged adjacent to the lesion (ROI1-ROI4, see Supplemental Figure 5). In Exp4NT, plants were imaged with 5 x and 20 x objective, frame average was set to 3 and z-stack size was adjusted to 15 steps (0.8 – 1.0 μm per step). Two consecutive ROIs were imaged adjacent to the lesion (ROI1 and ROI2, see Figure 5a). In Exp6NTNahG, plants were imaged with 20x objective, frame average was set to 3 and z-stack size was adjusted to 15 steps to cover at least 50 μm of the mesophyll. Two consecutive ROIs were imaged adjacent to the lesion (ROI1 and ROI2, see Figure 5a). See Supplemental Data Set 4 for zoom factor, digital zoom, optical zoom and for the number of analysed plants, leaves, lesions and ROIs for each experiment for each treatment and time point.

### Image Analysis

For the detection of chloroplast redox state, maximum projections from Z-stacks for each of three channels: chlorophyll fluorescence, roGFP fluorescence after excitation with 405 nm laser line and GFP fluorescence after excitation with 488 laser line were for each ROI exported from Leica LAS X software as a tif file.

Image analysis was performed using an in-house Matlab script. The images were analyzed with the following steps: import of tif files, conversion to grayscale, filtering out pixels of low intensity, conversion to binary format using spatial adaptive thresholding, a round or erosion and dilation to remove single pixel noise around the chloroplasts, followed by size-based segmentation of individual chloroplasts. The ratios of fluorescence intensities 405/488 were then calculated for each pixel belonging to the chloroplast masks obtained in the previous step. Results were calculated per image (normalized to the fraction of pixels belonging to chloroplasts) and per individual chloroplast in each image (see 405_488 in Supplemental Data Set 1 for 405/488 ratios).

In the next step, ROI1 and ROI2 were further divided into 5 regions (Bins; see Figure 3a), depending on the distance from the lesion (Bin 1 being closest, while Bin 5 being farthest from the lesion). Again, the 405/488 ratios were determined for each pixel inside chloroplast masks, then the first four Bins were normalized to the 405/488 ratio in Bin5, which was set to 1 (Supplemental Data Set 3).

For counting stromules, maximum projections from Z-stacks for each of two channels (chlorophyll and 488) were for each ROI exported from Leica LAS X software as a tif file. The stromules were counted manually on the 488 tif images, while the number of the chloroplasts was determined as described above. The number of stromules was normalised to the number of chloroplasts for the corresponding ROI (see Supplemental Data Set 4).

Details of all the parameters used in the image analysis can be found in the analysis scripts, which are available on https://github.com/NIB-SI/SensorPlantAnalysis.

### Statistics

To determine which factors contributed most to the variability in measured 405/488 ratio values, we used a linear mixed effects model (LME):

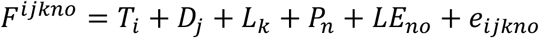

Where F is the fluorescence (405/488 ratio) measured on day *j* after treatment/ROI *i*, on leaf *o* of plant n belonging to plant line *k*; T_i_ is the mean (fixed) effect of treatment/ROI i; D_j_ is the mean (fixed) effect of time after treatment/viral infection j; L_k_ is the mean (fixed) effect of the specific plant line k; P_n_ is the (random) effect of plant n, assumed to be normally distributed with mean 0; LE_no_ is the (random) effect of the chosen leaf n on plant o, assumed to be normally distributed with mean 0; and e_ijkno_ is the residual error, assumed to be normally distributed with mean 0. If factors were not relevant for individual experiments (e.g. experiments run on a single line, or measured on a single day after treatment), they were removed from the mixed effects model. The final LME model was fitted with maximum likelihood to model the 405/488 ratio as the dependent variable, treatment/ROI, day post inoculation (dpi), transgenic line/genotype as fixed effects and plant and leaf as random effects.

Statistical analysis was implemented as an R based script, with significance calculation of the fixed factors performed using *anova* function (Satterthwaite’s method) from the package *lmertest* (Kuznetsova et al., 2017) and posthoc pairwise comparisons within levels of significant fixed factors performed using the *emmeans* function (Kendall-Roger method) available from the package *emmeans* (Lenth et al. 2018). The posthoc pairwise comparisons were performed between levels of individual fixed factors (eg, between ROI1 and ROI2 after viral infection), with and without stratification by the other fixed factors. For instance, comparison between ROI1 and ROI2 after viral infection, were performed for the whole experiment, as well as for the data available only for a single line, on a single day post infection, and for all combinations between line and day post infection (see Supplemental Data Set 2). All scripts were deposited to Github (https://github.com/NIB-SI/SensorPlantAnalysis).

## Supplemental Data

**Supplemental Data Set 1**: Phenodata and 405/488 ratios for chloroplastic redox detection experiments.

**Supplemental Data Set 2**: Statistical evaluation of observed relative redox changes in different conditions in individual experiments.

**Supplemental Data Set 3**: 405/488 ratios in five Bins of ROI1 and ROI2.

**Supplemental Data Set 4**: Phenodata and normalised number of stromules for stromules counting experiments.

**Supplemental Data Set 5**: Statistical evaluation stromule counts in different conditions in individual experiments.

**Supplemental Figure 1**: Relative chloroplast redox state after PVY and MOCK inoculation and H_2_O_2_ and DTT treatments.

**Supplemental Figure 2**: Individual cells with chloroplasts in oxidized redox state in ROI2 in PVY-inoculated redox state sensor plants.

**Supplemental Figure 3**: Detailed spatial analysis of relative chloroplast redox state around the cell death zone.

**Supplemental Figure 4**: Stromules in cell death zone proximity in PVY-inoculated pt-roGFP and pt-roGFP-NahG plants 5 days post PVY-inoculation.

**Supplemental Figure 5**: Stromule formation around the cell death zone is differently spatiotemporally regulated between potato genotypes.

**Supplemental Figure 6**: Relative chloroplastic redox state after treatments with a reference oxidant and a reductant in redox state sensor plants.

**Supplemental Method 1**: Stable Transformation

**Supplemental Method 2**: Construction of PVY-N605(123)-GFP infectious clone

## Acknowledgements

We thank Dr Fabrizio Cillo (Institute for Sustainable Plant Protection, Italy), for kindly providing the PVY-N605(123) plasmid, Prof. Andrej Blejec for help with statistical analysis and Lidija Matičič, Katja Stare, Barbara Dušak and Matej Rebek for technical assistance.

## Author contributions

KG and TL designed the research. TMP, MJ, TL performed the research. TL, KG, AŽ, TMP, ŠB analyzed the data. TL, KG, AŽ, TMP, MJ, ŠB, JB contributed to the writing or revision of the article.

